# Application-dependent assessment of the human exposure potential to microplastics

**DOI:** 10.1101/2025.09.29.679164

**Authors:** Michelle Klein, Bernd Giese

**Author notes:** Correspondence, Institute of Packaging and Resource Management, Department Applied Life Sciences, Hochschule Campus Wien, University of Applied Sciences, 1100 Vienna, Austria.

## Abstract

The presence of microplastic in the environment has become a matter of significant concern regarding its impact on the food chain and, moreover, human health. The potential ways in which humans may be exposed to microplastic include ingestion, inhalation, or dermal contact. To facilitate initial estimates of the potential for human exposure, a model has been developed that incorporates all of the major stages in the transfer of microplastic from its sources to direct human contact. Due to the scarcity of data available, a simplified gradation of exposure probability has been applied. Building on the results of published mass flow models for microplastics, the model is based on normalized values for the release of microplastics from different types of applications into environmental compartments, also considering the degradation of macroplastic. Published data investigating the contamination of different foods and beverages are evaluated, and the inhalation probability is assessed using air pollution data and breathing rates. In the final step, the potential for resorption through the gastrointestinal and the respiratory tract is estimated. The results obtained from this modelling indicate a high exposure potential for microplastic from tire wear via outdoor air and for various PET applications via indoor air. Furthermore, the results demonstrate a high potential for exposure via ingestion of food for plastics used in agriculture. The model’s results represent an initial attempt to estimate exposure probabilities for humans across all application categories of plastics and the most common polymers, taking into account the uncertainty of current research.

## 1. Introduction

Plastics are suitable for a wide range of applications due to numerous beneficial properties such as water resistance, low costs, high strength, low weight, and easy manufacturing. The versatile application possibilities led to a rising number of different plastic products. However, problems regarding their end-of-life or application-related emissions are often neglected (Andrady and Neal 2009). While their characteristics make plastic valuable during use, they tend to cause negative effects once released into the environment. Plastics’ durability results in slow degradation, taking up to 2500 years depending on the type and environmental conditions (Chamas et al., 2020). Their persistence and spatial distribution increase risks to ecosystems (Leslie et al., 2022; Ragusa et al., 2022). Plastics are omnipresent across various environmental areas, including river systems, polar regions, and mountain glaciers (Bergmann et al., 2022; Napper et al., 2020; Van Emmerik and Schwarz, 2020). Measurements indicate an exponential increase in plastic concentrations in sediments and oceans over the last decades (Lebreton et al., 2018; Martin et al., 2017). And it is expected that they will continue to increase as plastics production rises (Borrelle et al., 2020). Although subject to great uncertainties, the major sources of plastic pollution are most probably mismanaged macroplastics and tire wear particles (Sieber et al., 2020; Kawecki and Nowack, 2019), originating from littering, dumping, agricultural and construction waste, and improper disposal of consumer goods (Kawecki and Nowack, 2019).

The term microplastic (MP), introduced by Thompson et al. (2004), refers to plastic particles smaller than 5 mm, while nanoplastic refers to particles smaller than 1 µm, formed through abrasion, ultraviolet irradiation, hydrolysis, or biodegradation (Gigault et al., 2018; Mitrano and Wohlleben, 2020). Common sources of MP include tire abrasion, pre-production plastic pellets, paints, synthetic textiles, packaging, personal care products, and electronic equipment (Fendall and Sewell, 2009; Andrady, 2011; Barboza et al., 2018).

MPs can enter the human body through ingestion, inhalation, and skin contact, and have been detected in human thrombi, breast milk, newborn infants’ faeces, and have been shown to cross the blood-brain barrier (Ramsperger et al. 2023; Li et al., 2020; Zhang et al., 2021; Ragusa et al., 2022; Wu et al., 2022; Kopatz et al., 2023). While research has focused on the effects of MPs on the environment, knowledge of human exposure, tissue fate, and resulting health effects is limited (Ramsperger et al., 2023; Zhang et al., 2020). A holistic approach, which considers MP production, transmission, distribution, degradation, consumption, and its effects on human health, is so far still missing.

This work aims to establish a method for an initial rough estimate of human exposure to various plastic applications through ingestion, inhalation, and skin contact. Initially, relevant plastic applications and polymers were categorized for MP emissions. The normalized mass flow of MP emissions from each application into the environmental compartments water, air, and soil has been modelled, using data provided by published mass flow models for MP. Environmental macroplastic emissions were also considered, integrating degradation factors. To classify the relative exposure as high, medium, or low, limit values for the exact definition needed to be established, based on plastic production data in the EU. The model considers the MP content of a diverse range of foods and beverages. The transfer of MP into the human body is included as the last part of the model, although there is not yet much information on this in the literature.

## 2. Material and Methods

Excel 2013 was used for data processing, while the normalized model of plastic flows into environmental compartments was built in the graphic dynamic modelling program STELLA Professional©. The model was constructed using three main components: Stocks, Flows, and Action Connectors. Stocks store inflows and release them when outflows occur, while Flows fill or drain Stocks. Action Connectors facilitate interaction between Stocks and Flows. Within STELLA, the normalized flows from plastic applications to the environmental compartments were modelled.

Normalized models for MP emissions were constructed based on transfer coefficient data from Kawecki et al. (2018) and Kawecki and Nowack (2019). Separate models were built for seven polymers, each tailored to their specific transfer coefficients and plastic applications. Similarly, a macroplastic model was created, integrating degradation data and built separately for each polymer. A dedicated model for SBR emissions, including tire wear and artificial turfs, was developed using transfer coefficient data from Sieber et al. (2020).

To assess the human exposure potential starting from categories of plastic applications, three steps are evaluated and rated (Figure 1): 1) the probability of MP transfer in the environmental compartments air, soil and water, 2) the direct human availability (human external exposure) and 3) the possibility of transfer across tissue barriers, resulting in human internal exposure. At the end of each step, the obtained exposure probabilities are assigned to grades of exposure probability. To account for the current uncertainties in scientific literature, a simplifying estimation in the way of an allocation to three grades of probability was chosen for the model. Further differentiation would suggest precision, which is currently not available due to knowledge gaps, uncertainties, and limited comparability of published analyses.

**Figure 1.**
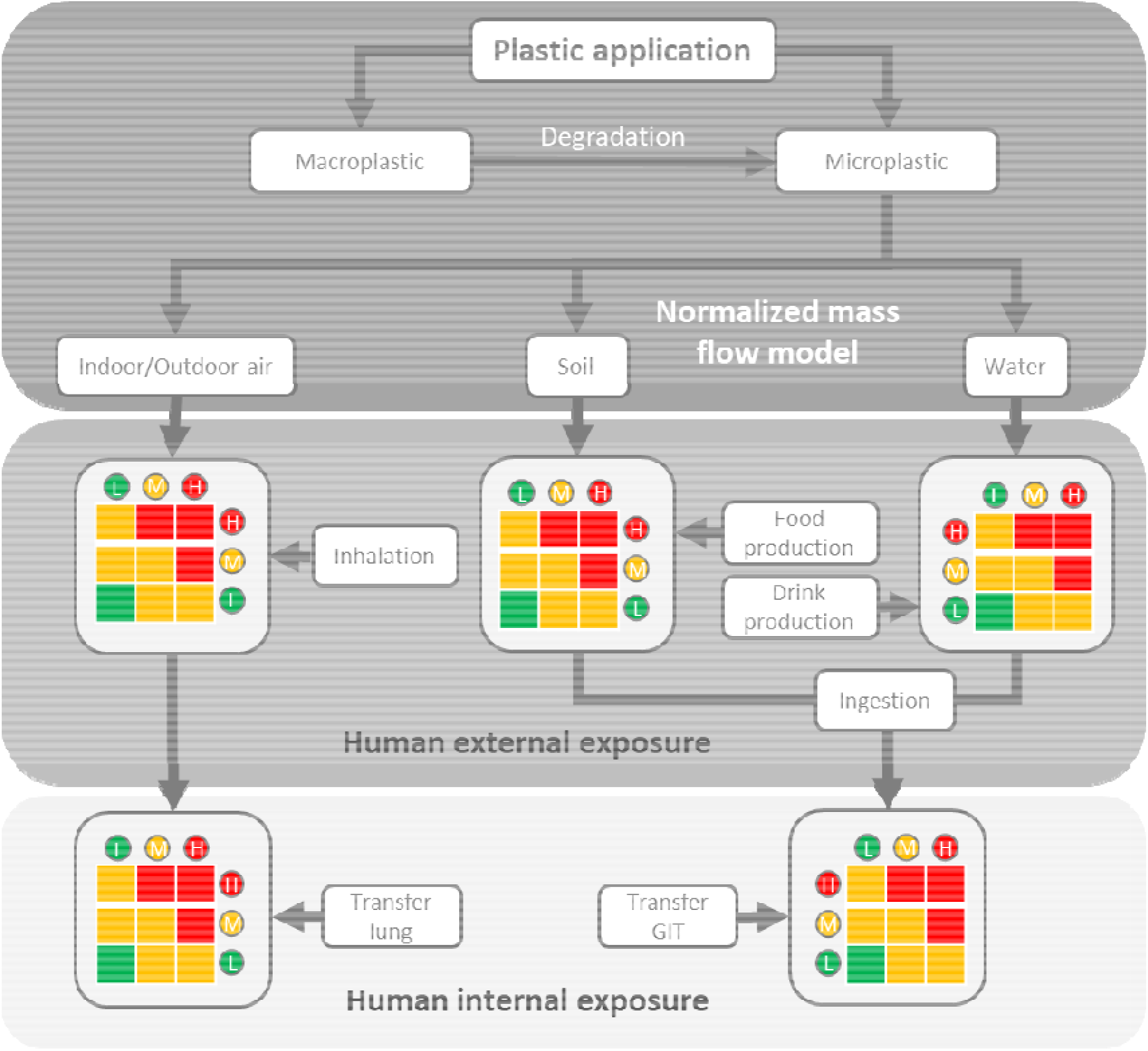
Overview of the modelling steps to estimate the human exposure to MP. L (green) = low exposure, M (yellow) = medium exposure, H (red) = high exposure, GIT = gastro-intestinal tract.

In the initial step, the normalized flow of MP shares from plastic applications released into the indoor and outdoor air, soil, and surface water and the release and degradation of macroplastic in these environmental compartments was modelled. The share of MP from the overall amount of material used for the respective applications is considered as a measure of the probability of occurrence in the environmental compartments, taking into account release and transfer rates as well as the degradation of macroplastic.

The external human exposure potential (step 2) is estimated by considering the probability of transfer to environmental compartments with a) the rated contaminations of food and beverages which are expected to be mainly caused by production processes and b) rates of ingestion and inhalation. For the respective combinations, we applied a precautionary scheme in which a combination of medium and low already leads to a medium instead of a low rating (Figure 1). MPs in soil were considered relevant for the potential uptake by plants. MPs in surface water were considered relevant for fish, mussels, salt, and drinking water. The following types of food and beverages considered in the model are not set in ratio to the potential MP shares in the respective environmental compartments due to their rather indirect relationships: honey, milk, processed foods and beverages such as sliced and packed meat, canned fish, beer, and soft drinks. For the exposure by inhalation, the intake into the respiratory tract had to be estimated considering breathing rates and air contamination with MPs (see supplementary information, chapter 4). Exposure by ingestion was determined by taking into account average consumption (see supplementary information, chapter 3).

In the last part of the assessment (step 3), the rating for the external human exposure potential is combined with the possibility of transfer across tissue barriers of the lung and the gastrointestinal tract (GIT) to determine the internal human exposure potential (Figure 1). The same combinatorial matrix as in step 2 is used for these estimates.

### 2.1 Selection of Plastic Applications

For the first step in the exposure assessment, the probability of MP transfer in environmental compartments, data from 41 plastic applications were obtained from Kawecki et al. (2018), Kawecki and Nowack (2019), and Sieber et al. (2020). Sieber et al. (2020) provide transfer coefficients for MP of tire rubber, while Kawecki and Nowack (2019) provide transfer coefficients for MP and macroplastic of LDPE, HDPE, PET, PP, PVC, PS, and EPS. These data are based on a former publication by Kawecki et al. (2018), which includes estimates for the annually used mass of these plastics for different applications. To decouple the shares of plastics considered in the current model from the used masses and still enable comparison between different applications, their mass per application category was normalized to the overall mass of plastics used per year.

### 2.2 Land-cover Data and Sludge Application

The models used to estimate the share of plastics in environmental compartments referenced Switzerland (Sieber et al. 2021; Kawecki and Nowack 2019). Therefore, adjustments were made in the normalized model to align calculations with EU conditions based on land-cover data from Eurostat (2021). For instance, the EU has 3.2% surface water compared to Switzerland’s 4.3%. Similarly, residential soil accounts for 4.2% in the EU compared to 7.5% in Switzerland. Agricultural soil in the EU is 24.2% compared to Switzerland’s 35.9%, while natural soil is estimated at 68.3% for the EU and 52.3% for Switzerland.

Another adjustment was made to the sludge application. While Switzerland prohibits its use on agricultural land, most EU countries allow it as fertilizer. Based on 2015 data, the average share of sludge application across EU Member states is 29.27%. This percentage was used as the transfer coefficient from sludge to agricultural soil in the model, reflecting its common use on farmland (Roskosch et al. 2018).

### 2.3 Assumptions

One assumption in this model is the absence of previous emissions, focusing solely on current data inputs to estimate human exposure probabilities. For categories lacking European data, Swiss figures were scaled up using a 1:60 population ratio assumption as applied in Kawecki and Nowack (2019). This scaling was applied to categories such as disposable cleaning cloths, cotton swabs, tampons, tampon applicators, wet wipes, panty liners, and sanitary napkins. Data on SBR production for tires and artificial turfs were not available, therefore careful estimates were made. Using data from Verschoor et al. (2016), the SBR content of tires was calculated as 24.3% of the total tire weight, summing up to 2.14 kg SBR per tire. Based on vehicle registration data from the World Health Organisation (WHO) (2020) and assuming four tires per vehicle, SBR production in Europe was estimated at 3,053,978 kt. For artificial turfs, infill amounts were taken from Hann et al. (2018), yielding an estimate of 1.8 million tonnes of SBR used across 51,616 fields in Europe.

### 2.4 Integration of Degradation of Macroplastic

In the first step, MP data from Kawecki and Nowack (2019) and Sieber et al. (2020) including the modifications described above have been used to calculate MP shares in environmental compartments. Macroplastic emissions under consideration of degradation factors taken from Schwarz et al. (2023) have been modelled in addition to also considering MP emergence from macroplastic degradation. The data of Schwarz et al. (2023) provide degradation rates for eight polymers. These rates, applied to the macroplastic model, yield the share of secondary MP which is finally added to the share of primary MP in the normalized model. The degradation timeframe of one year in the normalized model corresponds with the assumption in Schwarz et al. (2023). Table 1 presents degradation rates for the eight polymers, with values based on data presented by Schwarz et al. (2023) and assumptions for PET degradation in agricultural soil made by Schwarz et al. (2023) (marked by an asterisk). Environmental compartments considered in this study include indoor air, outdoor air, surface water, and agricultural soil, following studies by Kawecki and Nowack (2019), Sieber et al. (2020), and Schwarz et al. (2023).

**Table 1.**
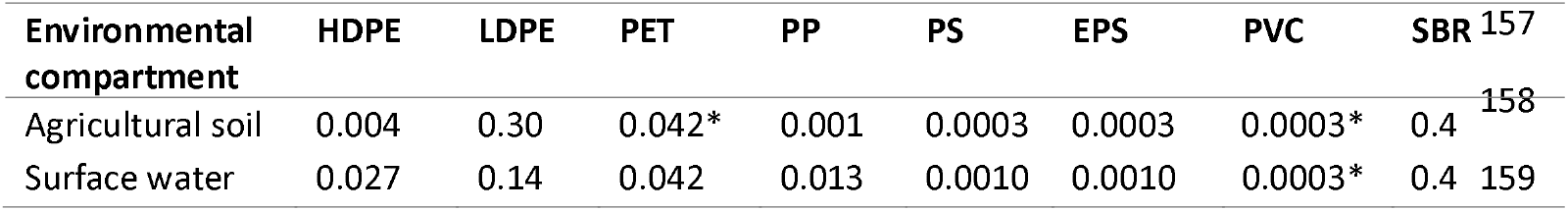
Degradation coefficients in percentiles for eight polymers in environmental compartments in the timeframe of a year. Values marked with an asterisk are assumptions of Schwarz et al. (2023).

### 2.5 Limit Values

To account for insecurities and data gaps, the exposure potential in all intermediate steps and the final result of the model for human exposure is expressed using three levels: high, medium, and low. To assign the shares of MP in environmental compartments calculated in the first step to these three levels, limit values needed to be set. The limits are oriented at the scale of the produced mass of polymer in kt in Europe in 2014 for the respective application (Kawecki and Nowack 2019) (Table 2).

**Table 2.** Limit values for exposure classification are based on the mass of produced polymer per plastic application per year in kt.

| kt/year | Low | Medium | High |
| --- | --- | --- | --- |
| ≤10 | <1% | 1 – 10% | >10% |
| 10 – 100 | <0.1% | 0.1 – 1% | >1% |
| 100 – 1,000 | <0.01% | 0.01 – 0.1% | >0.1% |
| 1,000 – 10,000 | <0.001% | 0.001 – 0.01% | >0.01% |

As for the share of MP in environmental compartments, limit values are defined to rate the potential uptake as low (≤ 100), medium (100 – 1,000), and high (≥ 1,000), with reference to the number of annually ingested/inhaled particles. The same values are chosen for the uptake via inhalation. Due to the lack of toxicological data, these limits do not reflect any effect thresholds. They merely represent a classification chosen by the author.

## 3. Results

### 3.1 Model Building

For the first step, to model emission probabilities from plastic application to environmental compartments, data provided in the Supporting Information by Sieber et al. (2020) and Kawecki and Nowack (2019) were used. Two application categories were provided by Sieber et al. (2020), namely “Tire wear” and “Artificial Turfs”. Kawecki defined 57 different application categories of which 41 were used for this work. Pre-consumer and post-consumer processes – which included primary and secondary production, textile and non-textile manufacturing, transport, and several recycling categories – are not included in this work, as production and end-of-life processes are not considered under the term “plastic application”.

In the next step, categories defined by Kawecki et al. were merged for the modelling process in STELLA, when their transfer coefficients were comparable and when the categories could be considered to belong to the same general field of application. This simplification resulted in a further reduction of categories from 39 to 23 (plus two categories based on data provided by Sieber et al. (2020)).

### 3.2 Emissions from Applications to Environmental Compartments

#### 3.2.1 Indoor Air and Outdoor Air

The environmental compartment “air” is divided into indoor and outdoor air. Outdoor emissions, like tire wear, are only considered for outdoor air. For emissions happening indoors, e.g. “household plastics”, an exposure potential in indoor air and outdoor air due to air ventilation is assumed in the model.

“Household plastics” made of HDPE and PS and coverings made of PVC are rated with a medium exposure potential for emissions to indoor air. PP household plastics yield the same MP release to indoor air but are rated with high exposure potential due to their ten times higher production input (Kawecki and Nowack 2019) (Table 3). Consistent values for the release to indoor air across different applications and polymers are due to unspecific transfer coefficients (Kawecki and Nowack 2019).

**Table 3.** Plastic applications and the specific polymer with emissions to indoor and outdoor air. Limit values are applied based on the production input and the released portion of the input. The production input in kt is based on data provided by Kawecki and Nowack (2019).

|  | Plastic Application | Polymer | Production input<br>in kt | Applied limit<br>value | Input ending up<br>in air in % |
| --- | --- | --- | --- | --- | --- |
| Indoor Air | Household plastics | HDPE | 130 | 0.01 – 0.1%<br>(medium) | 0.000211 |
|  | Household plastics | PS | 110 | 0.01 – 0.1%<br>(medium) | 0.000211 |
|  | Household plastics | PP | 1,300 | >0.01% (high) | 0.000211 |
|  | Coverings | PVC | 500 | 0.01 – 0.1%<br>(medium) | 0.000211 |
| Outdoor Air | Household plastics | HDPE | 130 | <0.01% (low) | 0.00421 |
|  | Household plastics | PP | 1,300 | <0.01% (low) | 0.00421 |
|  | Household plastics | PS | 1,300 | <0.01% (low) | 0.00421 |
|  | Coverings | PVC | 500 | <0.01% (low) | 0.00421 |
|  | Insulation | PS | 90 | <0.01% (low) | 0.0116 |
|  | Insulation | EPS | 1,100 | 0.001 – 0.01%<br>(medium) | 0.00211 |
|  | Tires | SBR | 3,053,978 | >0.01% (high) | 0.063 |

Few plastic applications contribute to outdoor air emissions, including “insulation” made of PS and EPS, “household plastics,” “coverings,” and SBR from “tire wear”. Emissions from indoor air to outdoor air, as assumed by Kawecki and Nowack (2019), are diluted, resulting in a rating as “low” exposure potential for “household plastics” made of HDPE, PP, and PS and “coverings” made of PVC (Table 3). “Insulation” made of PS emits 0.0116% of the total produced mass to outdoor air and is rated low, while EPS with 0.00211% ending up in outdoor air is rated medium due to the higher production input of EPS.

#### 3.2.2 Water

In comparison to the results for indoor and outdoor air, more plastic applications lead to an exposure probability that can be classified as high for surface water (Table 4). Looking at PET, the category “consumer bags” showed a medium rating, while the other consumer categories “films”, “bottles” and “other consumer packaging” showed a high potential. This might be due to the high littering rate assumed by Kawecki and Nowack (2019).

**Table 4.** Plastic applications and the specific polymer with emissions to water. Limit values are applied based on the production input and the released portion of the input.

| Plastic Application | Polymer | Production input<br>in kt | Applied limit value | Input ending up<br>in water in % |
| --- | --- | --- | --- | --- |
| Consumer films | PET | 230 | >0.01% (high) | 0.000457 |
| Consumer films | PP | 600 | >0.01% (high) | 0.152 |
| Consumer films | PVC | 26 | 0.1 – 1% (medium) | 0.161 |
| Consumer bags | PET | 900 | 0.01-0.1% (medium) | 0.000387 |
| Consumer bags | PP | 330 | >0.1% (high) | 0.142 |
| Consumer bags | PVC | 5 | <1% (low) | 0.166 |
| Consumer bags | PS | 10 | <1% (low) | 0.179 |
| Consumer bottles | PET | 1,300 | >0.001% (high) | 0.00000890 |
| Consumer bottles | PP | 33 | <0.1% (low) | 0.00445 |
| Consumer bottles | PVC | 10 | 0.1 – 1% (medium) | 0.166 |
| Other consumer packaging | PET | 280 | >0.01% (high) | 0.000193 |
| Other consumer packaging | PP | 1,200 | >0.01% (high) | 0.0436 |
| Other consumer packaging | PVC | 200 | >0.1% (high) | 0.224 |
| Other consumer packaging | PS | 600 | 0.01 – 0.1% (medium) | 0.0476 |
| Other consumer packaging | EPS | 290 | <0.01% (low) | 0.000271 |
| Household plastics | PET | 130 | <0.01% (low) | 0.00183 |
| Household plastics | PP | 1,300 | 0.001 – 0.01% (medium) | 0.00183 |
| Household plastics | PS | 110 | <0.01% (low) | 0.00183 |
| Fabric coating | PVC | 110 | >0.1% (high) | 0.15 |
| Insulation | PS | 90 | <0.1% (low) | 0.0469 |
| Insulation | EPS | 1,100 | >0.01% (high) | 0.0448 |

#### 3.2.3 Agricultural Soil

Kawecki and Nowack (2020) classify soil into three types: residential, natural, and agricultural. In the present model, agricultural soil, used for food production, is directly linked to MP occurrence in the food chain, while other soil types are not. Especially for plastic applications in an agricultural context high transfer rates to agricultural soil are assumed. These applications are mainly classified as having a high exposure potential (Table 5).

**Table 5.** Plastic applications and the specific polymer with emissions to agricultural soil. Limit values are applied based on the production input and the released portion of the input.

| Plastic Application | Polymer | Production input in kt | Applied limit value | Input ending up in agricultural soil in % |
| --- | --- | --- | --- | --- |
| **Agricultural packaging films** | LDPE | 190 | >0.1% (high) | 1.77 |
| **Agricultural films** | LDPE | 900 | >0.1% (high) | 3.11 |
| **Agricultural films** | PVC | 30 | >1% (high) | 2.01 |
| **Agricultural pipes** | LDPE | 180 | >0.1% (high) | 1.24 |
| **Agricultural pipes** | PVC | 40 | 0.1 – 1% (medium) | 0.111 |
| **Agricultural other** | LDPE | 140 | >0.1% (high) | 1.24 |
| **Agricultural other** | PVC | 25 | 0.1 – 1% (medium) | 0.111 |
| **Agrotexiles** | PET | 9 | <1% (low) | 0.684 |
| **Agrotexiles** | PP | 260 | >0.1% (high) | 0.659 |
| **PCP** | HDPE | 4 | >10% (high) | 22.3 |
| **PCP** | PET | 4 | >10% (high) | 22.3 |
| **PCP** | PP | 0,009 | >10% (high) | 22.3 |
| **Household plastics** | PP | 1,300 | 0.001 – 0.01% (medium) | 0.00912 |
| **Fabric coatings** | PVC | 110 | 0.01 – 0.1% (medium) | 0.0901 |
| **Insulation** | EPS | 1,100 | 0.001 – 0.01% (medium) | 0.00242 |
| **Tires** | SBR | 3,053,978 | >0.01% (high) | 0.198 |

The majority of MPs are derived from the degradation processes of macroplastic. For agricultural packaging films, 1.77% of these films end up as MPs from degraded macroplastic in agricultural soil, while for agricultural films, 2% derive from primary microplastic and 1.11% through degradation. For agricultural pipes and the category “agricultural other”, 0.1% of input are primary microplastic and 1.14% from degraded macroplastic. This finding indicates that, with the exception of “agricultural films”, the majority of microplastic particles entering agricultural soil during the specified period are attributable to degraded macroplastic.

Few plastic applications beyond agriculture exhibit medium or high exposure potential. For instance, PET in “wet wipes” poses a medium exposure risk due to the high production volume and the likelihood of disposal via flushing, leading to eventual deposition in agricultural soil through sewage sludge and later application as fertilizer on agricultural soil. Similarly, “fabric coating” made of PVC shows a high exposure potential in agricultural soil due to similar pathways.

Another medium exposure potential is attributed to “household plastics” made of PP. Since this application emits MPs into the air, their presence in agricultural soil may result from atmospheric deposition.

### 3.3 Human Exposure through Ingestion

To date, contamination of various foods and beverages by MPs has been reported, including honey (Liebezeit 2015), meat (Kedzierski et al. 2002, Habib et al. 2022), fish (Gündoli⍰du and Köşker 2023), sugar (Liebezeit 2013), salt (Johnson et al. 2020), tap and bottled water (Oßmann et al. 2018) and beer (Diaz-Basantes et al. 2020). There are several steps in the food chain, where contamination with MP can occur. This includes processing, treatments, packaging, and distribution (Toussaint et al. 2019).

For the direct human availability of MP, data from studies on MP in different foods and beverages are used. Values for different polymer types are evaluated whenever found in the respective studies. Based on these data in combination with annual consumption rates of the respective foods, the annual uptake of MPs is calculated (see supporting information).

When comparing the results for the different foods, the annual uptake by humans ranges from less than one MP in rice to up to 398,520 in tomatoes (Table 6). High exposure probabilities were identified for sliced meat, fruits, and vegetables as well as types of salt and sugar.

**Table 6.**
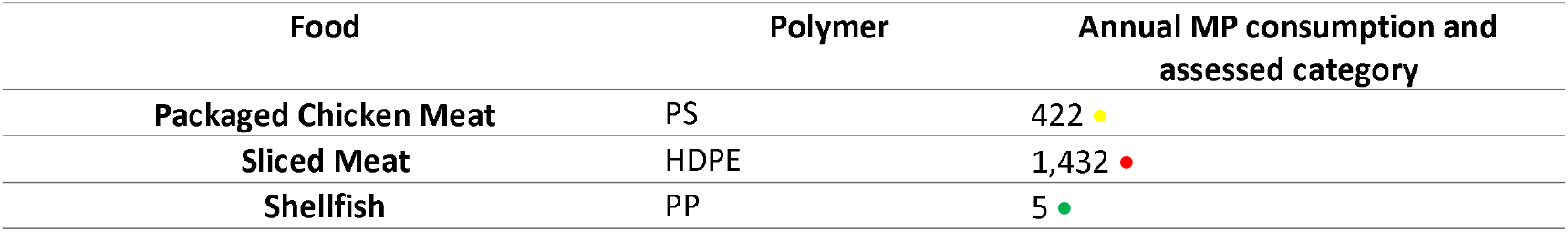
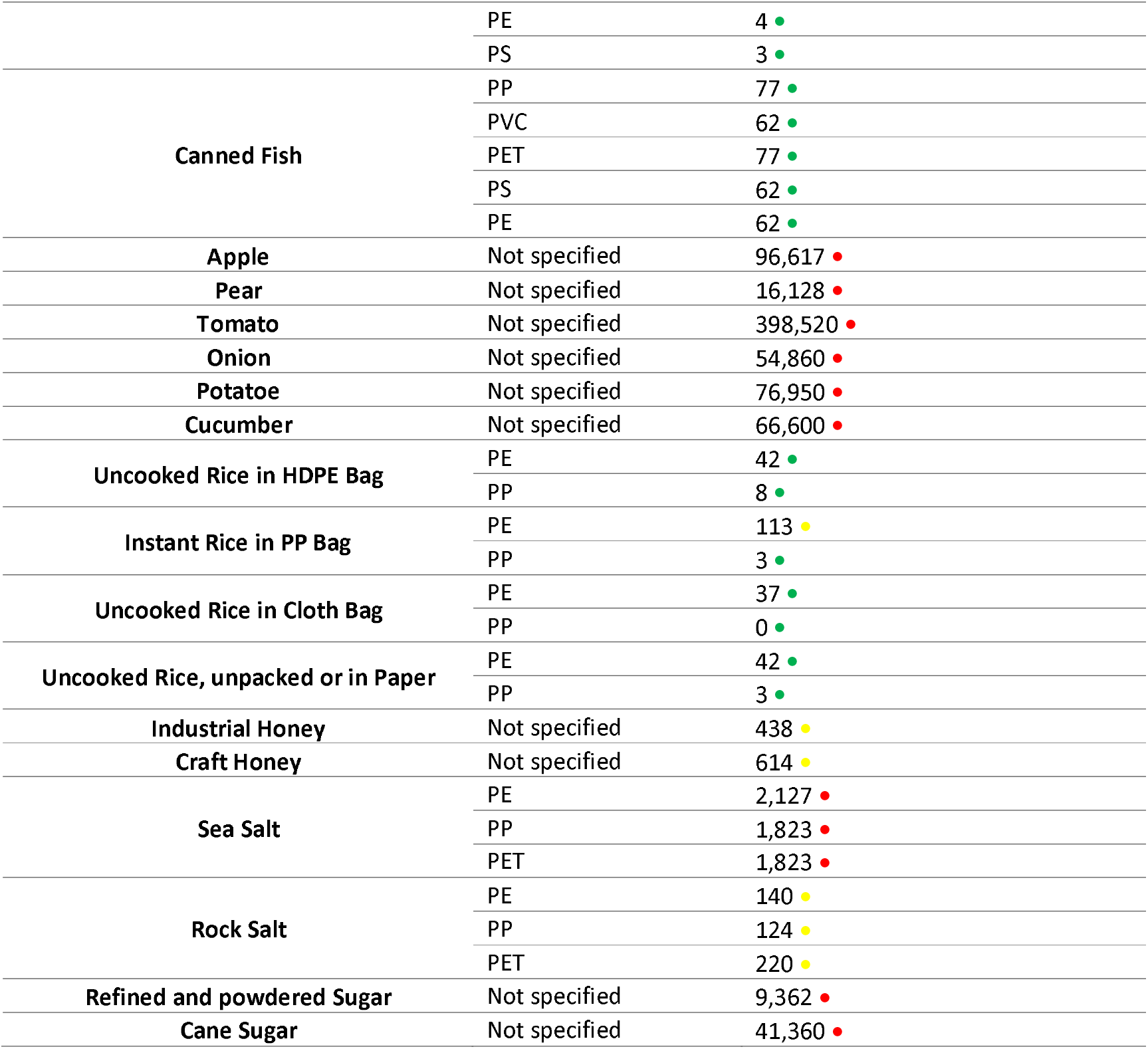
Overview of foods and annually consumed MPs by polymer and the respective exposure rating indicated by the coloured dot (low – green, medium – yellow, high – red).

The high content of MPs in sliced meat is most probably due to the wearing of the cutting boards (Habib et al. 2022). Therefore, the contamination can be traced back to the meat processing practices and not to the potential uptake of MPs by the animal and translocation into edible parts. Fruits and vegetables show the highest contamination rates of all analysed foods. As for those products, only few data were available, it would be necessary to conduct more studies to back up the data and to evaluate the origin of the contamination. As fruits and vegetables were peeled, air deposition of MPs during transport, storage, and packaging can be excluded. Therefore, contamination could be due to the uptake of MPs during the growth period, eventually from the soil.

The contamination in sea salt can be reasoned with high plastic pollution in the marine environment. For rock salt, a possible source of MP contamination could be air deposition. For sugar, sources would need to be identified. A potential uptake from the soil into sugar canes and sugar beets is possible. The contamination in honey can be traced back to improper handling and air deposition of MPs according to Liebezeit (2015).

In all cases, contamination during food processing needs to be considered.

The presence of PS particles from packaging in chicken meat indicates another issue for food safety. Food processing and packaging should avoid contamination, instead of adding to the problem. Few studies so far are available on the influence of MP from packaging in the food chain, which indicates a need to further investigate this issue.

Other foods received a low rating. For rice, the packaging seemed to have no significant influence on MP contamination, compared to the amounts found in packaged meat. Also, the contamination of fish and shellfish and the respective potential annual uptake of MPs is low, compared to the amounts taken up from sea salt, although both are influenced by marine plastic pollution. From the rice samples, the instant rice in PP packaging showed an elevated amount of PE, which was rated medium. As the packaging does not contain PP, the PE may stem from the processing of the rice.

No medium ratings were assigned for beverages (Table 7). Tap water is the only beverage with a rating of low exposure probability, which might be due to good filtration practices.

**Table 7.** Overview of beverages and annually consumed MPs by polymer and the respective exposure rating indicated by the coloured dot (low – green, medium – yellow, high – red).

| Beverage | Polymer | Annual MP consumption and assessed category |
| --- | --- | --- |
| Tap Water | PS | 0.004 ● |
|  | PET | 0.05 ● |
|  | PVC | 0.01 ● |
|  | PE | 0.005 ● |
| Water from single-use PET Bottle | PP | 29,218 ● |
|  | PE | 2,045 ● |
|  | PET | 260,044 ● |
| Water from old reusable PET Bottle | PP | 91,979 ● |
|  | PE | 49,669 ● |
|  | PET | 751,470 ● |
| Water from new reusable PET Bottle | PP | 29,660 ● |
|  | PE | 16,016 ● |
|  | PET | 242,320 ● |
| Water from Glass Bottle | PP | 159,622 ● |
|  | PE | 319,243 ● |
|  | PET | 24,984 ● |
| Bagged Tea | PET | 6,172,500,000,000 ● |
| Industrial Beer | Not specified | 2,970 ● |
| Craft Beer | Not specified | 2,022 ● |
| Skim Milk | Not specified | 2,268 ● |
| Soft drinks | Not specified | 2,950 ● |

The same consumption rates for tap water were applied to bottled water. Bottled water received consistently high ratings, showing the significance of potential contamination through filling processes and packaging materials. The contamination in old PET bottles compared to new bottles is three times higher. The PET content in glass bottles is ten times lower than in new reusable PET bottles, showing the significance of packaging, as glass bottles do not contain PET. On the other hand, the findings of PET in glass bottles indicate contamination during the bottling process.

The uptake of MPs through the consumption of beer, milk, and soft drinks lies within the same range. For beer, the selected packaging was glass, therefore contamination during filling is a potential source. For milk and soft drinks, the packaging options included beverage cartons and PET bottles, showing that contamination due to packaging material is a considerable option.

#### 3.3.1 Translocation through the Gastrointestinal Tract and Particle Uptake

Ingestion is the primary pathway for human exposure to MPs, due to constant consumption (Yee et al. 2021). MPs entering the gastrointestinal tract (GIT) become entrapped in mucus and may be transported to the epithelial layer (Hussain et al. 2001). Transcellular transport of particles below 100 nm occurs through endocytosis, while those larger than 100 nm are paracellularly transported through junctional complexes. Tight junctions are interrupted by goblet cells, which allow the paracellular transport of particles (Vancamelbeke and Vermeire 2017; Volkheimer 1977). The lamina propria beneath the epithelium contains immune cells, with M-cells transporting particles into pockets for internalization by dendritic cells. Those can internalize PS particles up to 15 μm in size (Hussain et al. 2001, Owen et al. 1999, Foged et al. 2005). Schwabl et al. (2019) found a mean of 28 MPs/g colon sample, primarily composed of 96% fibres with a length of 1 mm. Colon samples contained mainly fibres, while stool samples contained fragments and film-shaped particles. Given the large knowledge gaps regarding polymers other than PS and the assumption that especially the small fraction of particles represents the majority in the number-size distribution, the ability of MP to cross the intestinal tissue barrier is provisionally assessed as medium.

### 3.4 Human Exposure through Inhalation

#### 3.4.1 Indoor Air

Inhalation rates have been described by Abbasi (2021), which argued, that an adult inhales around 15 m^3^ air per day and spends 14 hours a day indoors, concluding, that within these 14 hours, 8.75 m^3^ air is inhaled.

Quantifications of fibres have been done extensively by Dris (2016). Dris (2016) measured 5.4 fibres/m^3^ indoor air, of which 33% turned out to be of polymeric origin, resulting in approx. 1.8 polymer fibres/m^3^. A first calculation shows, that within breathing 8.75 m^3^ indoor air daily, a total of about 15.6 polymer fibres is inhaled by an adult. Liu, Li et al. (2019) characterized those polymer fibres, showing that 80% are Polyester, and around 20% are of other origin. Since PET fibres are of interest for this model, the proportion of polyester is assumed for the PET content, which corresponds to approx. 12.5 PET fibres that are inhaled daily through indoor inhalation. Liu, Li et al. (2019) describe polymer particles in indoor air as of fibrous shape in 88% of the particles and non-fibrous in 12% of the particles. As the average adult consumes about 15.6 fibres per day, approx. 2.1 fragments are inhaled on average per day.

The polymer type of fragments in outdoor air was specified by Wright et al. (2020). As there was insufficient data available on fragments in indoor air, the same distribution as in outdoor air was considered. For this model, the relevant polymers are PP (12.5%), PE (12.5%), PVC (9.4%), PET (11.3%) and PS (18.9%). EPS was not listed specifically, but as 8.4% were considered as “others”, this value was used for EPS in the model (Liu, Li et al. (2019). For PE, no distinction was made between LDPE and HDPE. Therefore, 12.5% was taken for the proportion of LDPE and HDPE.

#### 3.4.2 Outdoor Air

For the quantification of fibres inhaled in outdoor air, data from Dris (2016) are again taken as a base for the calculation. According to Dris, 0.9 fibres are inhaled per m^3^. As reported by Abbasi (2021), the average adult spends 10 hours per day outdoors and inhales 15m^3^ per day, 6.25 m^3^ of outdoor air is inhaled daily. This results in an inhalation of approx. 5.6 polymer fibres per day. Wright et al. (2020) measured polymer content in outdoor air, reporting 18.8% of fibres specified as Polyethylene Terephthalate and Polyester (PET/PES). This results in a daily PET/PES fibre inhalation in outdoor air of approx. 1.1 fibres. Fragments in outdoor air have been characterized by Wright et al (2020). Liu, Li et al. (2019) describe that 73.7 % of polymer particles in outdoor air are fibrous and 26.3 % are non-fibrous. As the average adult consumes about 5.6 fibres per day, about 2 fragments are inhaled.

An assessment of tire and road wear particles in outdoor air was done by Panko et al. (2019). At the sampling sites in London, an average of 18.35 μg/m^3^ of PM2.5 particles were measured. The amount of PM10 particles summed up to 47.43 μg/m^3^. In a simplifying assumption, considering a spheric shape and the density of styrene-butadiene rubber (SBR) as a typical synthetic rubber of tires, the total number of tire rubber particles can be calculated from the given results (see Supporting Information). With 6.25m^3^ of daily inhaled outdoor air, a number of up to 57 PM10 and 14 PM2.5 SBR particles per day can be expected to be contained in the inhaled air.

As for food and beverages, data were scaled up to a time frame of one year. The same limit values as applied to ingested particles were considered (Table 6). In Table 8, data for the annually inhaled number of MPs are shown with a colour code depending on the categorization into a low, medium, and high exposure potential.

**Table 8.**
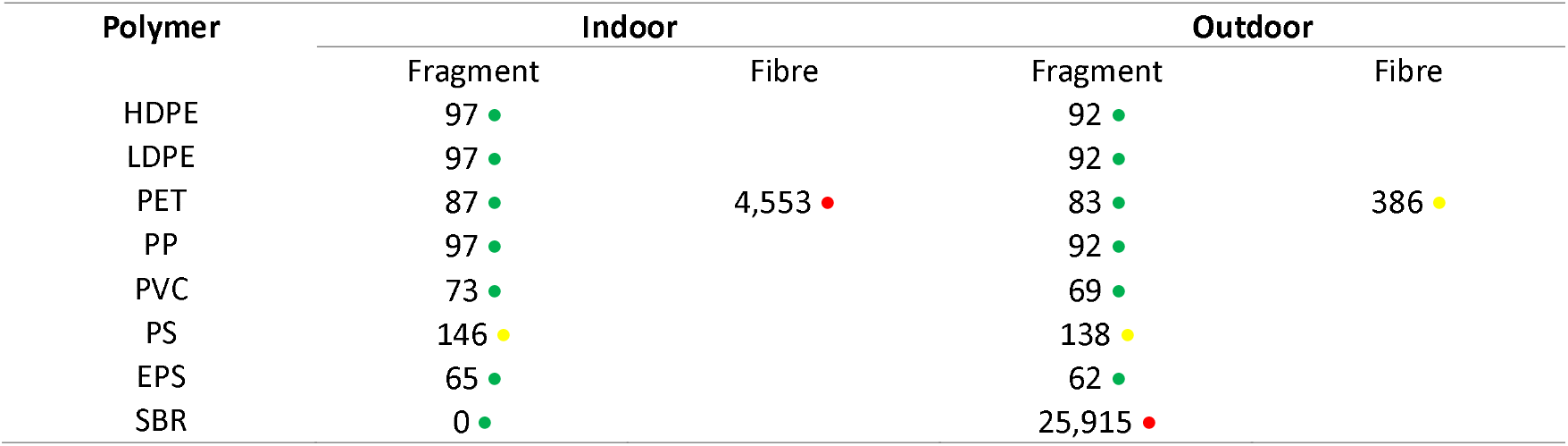
Estimated number of annually inhaled MPs. The colour code symbolizes the categorization into low (green), medium (yellow), and high (red) exposure.

A high exposure potential in indoor air is found for PET fibres, in outdoor air the exposure probability to SBR is rated as high. A medium assessment was received for PET fibres in outdoor air, which might be due to the shedding of textiles in outdoor areas or air ventilation from indoor environments. Also, PS fragments in indoor air and outdoor air received a medium rating.

#### 3.4.3 Translocation through the Lung

The body’s defense mechanisms against inhaled particles vary depending on their size. Larger particles, like PM10 (≤ 10 μm), are trapped in the nasopharyngeal area by mucus and hair, preventing further penetration. Fine particles, such as PM2.5 (0.1 – 2.5 μm), can reach the bronchioles and alveoli, while PM0.1 (up to 0.1 μm) can translocate across the alveolar epithelium (Kernchen et al 2022). The epithelial layer, with mucus produced by goblet cells, helps trap and remove inhaled particles through ciliary beating, allowing expulsion from the body. In this way, even PM0.1 can be caught up and discharged from the body (Schraufnagel 2020; Cooper and Loxham 2019). A recent review reports on microplastic particles in the human lung that are smaller than 500 μm. (Huang et al. 2022; Jenner et al. 2022 cited in Ramsperger et al. 2023). However, due to the still limited number of studies (Ramsperger et al. 2023), and regardless of possibly effective defense mechanisms, it is assumed that the probability of MP translocation through lung tissue is medium.

### 3.5 Human Exposure through Dermal Contact

The main product category relevant for skin exposure to MPs is personal care products (PCPs).

The translocation of MPs through intact skin is complex and limited. The outer skin layer, the stratum corneum, acts as an effective barrier against foreign matter and pathogens. One potential pathway for MPs to pass intact skin is through the trans appendageal pathway, involving hair follicles, sweat glands, or sebaceous glands. Although this pathway covers only a small percentage of the skin surface (0.1-1.3%), its relevance depends on MP concentrations in PCPs (Schneider et al., 2009; Desai et al., 2010). Particle size is a crucial factor for translocation through the skin, along with particle properties and skin health (Schneider et al. 2009). Studies have shown that negatively charged particles sized between 50 to 500 nm can penetrate the skin, especially when the skin is mechanically stressed (Kohli and Alpar, 2004). Larese Filon et al. (2015) suggest a maximum size of 4 nm for particles to cross intact human skin, while particles sized from 4 to 20 nm may translocate through both intact and damaged skin. Nanoparticles between 21 to 45 nm can pass through damaged but not intact skin, and particles larger than 45 nm are unable to translocate through either healthy or damaged skin. Limited research is available on the transfer of microplastic particles through the skin. For MP in PCPs, Ramsperger et al. (2023) consider ingestion and inhalation as the primary routes of exposure.

In conclusion, we currently assume that the skin forms a robust barrier against particulate matter, including MPs. Given that the most used polymers in PCPs are HDPE, PP, and PET, only these three polymers are assessed. The likelihood of MP translocation through human skin is therefore rated as low and was not taken into account in the further evaluation.

## 4. Discussion

This study aims to provide a simplified yet comprehensive assessment of human exposure to MPs emitted by various applications and transported through different environmental compartments. Seven common polymers (LDPE, HDPE, PET, PP, PVC, PS, and EPS) covering 57% of EU polymer production were selected for this analysis (PlasticsEurope 2023). Additionally, SBR elastomer was included due to its significant contribution from tire wear. Human exposure levels for each plastic application were categorized as low, medium, or high using a traffic light scheme, inspired by existing methodologies such as NanoRiskCat (Hansen et al. 2014) and a marine environment hazard ranking system (Yuan et al. 2022), which use a five-class-system. Due to data gaps, uncertainties, and only limited comparability of exposure and fate studies (cp. Ramsperger et al. 2023), a simplified approach with three exposure classes was adopted.

### 4.1 Potential of Transfer to Air, Water, and Agricultural Soil

This work focused on the human MP exposure potential of consumer applications of plastics. In future work improvements of the database of the presented model are necessary. Despite detailed calculations by Kawecki and Nowack (2019) for textile applications, generalized values are applied for environmental emissions. Additionally, textile emissions to agricultural soils are a significant concern (Zhang et al. 2021) and are overlooked due to nation-specific assumptions about the fate of sewage sludge (Kawecki and Nowack 2019). In this study, the transfer of plastic fibres from textiles to agricultural soils through the application of sewage sludge is considered. This enables a more realistic scenario, as the application of sewage sludge is still common practice in most European countries according to Roskosch et al. (2018).

The model reconstructs the transfer of HDPE, PET, and PP polymers to WWTPs, leading to MP emissions into surface water and likely transfer to agricultural soil via sewage sludge. Exposure probabilities for those sanitary applications have been rated as high for the transfer via drinks and medium for the transfer via foods. High MP transfer from agriculturally used plastics to soil aligns with the literature (Ren et al. 2023), especially considering macroplastic degradation as modelled in this study.

Consumer and non-consumer plastic applications which mainly emit to surface water, received mixed ratings depending on polymer type, with littering considered only in these categories in the model provided by Kawecki and Nowack (2019). Kawecki and Nowack (2019) data suggest medium to high transfer probabilities to surface water for building and construction materials. The transfer probabilities of HDPE, PET, and PP polymers for PCPs via WWTPs result in medium MP emissions into surface water and are highly likely to transfer to agricultural soil via sewage sludge. However, with the European Commission’s microplastics restriction, products containing MP, except MP in solid matrices, may no longer be placed on the market after October 2023, with some exceptions for products with transitional periods or stocks. High MP transfer from agriculturally used plastics into soil corresponds with existing literature about plastic mulch use on cropland, especially considering macroplastic degradation. The inclusion of sludge application and degradation coefficients for macroplastics in this model showed the relevance of the degradation of plastics in agriculture as a soil pollution problem and improved the understanding of the impact of PCP and sanitary articles in terms of emissions on agricultural soils and the food chain.

The existing model by Kawecki and Nowack (2019) primarily indicates low emissions of consumer and non-consumer plastic applications to agricultural soil, with littering considered mainly in these categories. However, oversight of potential surface water emissions underscores the need for model enhancements, as the transport of microplastic particles via erosion and wind is ignored. Due to the lack of available data, it was also not possible to add this pathway to this model; this is a topic for future research to be included in the modelling of microplastic exposure.

A separate part of our model assessed environmental emissions from tire wear and artificial turf, with tire wear identified as a significant emitter of MPs to outdoor air, soil, and water. Despite the model’s generally good performance in identifying typical MP emission sources, textiles require further refinement with more differentiated transfer coefficients.

### 4.2 Human Exposure through Ingestion and Inhalation

Several studies provide data on MP contamination of foods and beverages, revealing notable trends in contamination levels that impact human consumption (Liebezeit 2015; Kedzierski et al. 2002, Habib et al. 2022; Gündoli⍰du and Köşker 2023; Liebezeit 2013; Johnson et al. 2020; Oßmann et al. 2018; Diaz-Basantes et al. 2020; Toussaint et al. 2019; Aydin et al. 2023). Despite concerns over marine pollution, fish and shellfish show comparably low contamination, while rice seems to be less affected by plastic packaging compared to meat or bottled water. Limited research on MPs in fruits and vegetables underscores the need for further investigation into contamination levels and human exposure probabilities. Future studies should also further examine the effects of food processing and packaging on MP contamination. Inconsistent methodologies and level of detail and reported specifics hinder comparisons between studies. The estimated levels of human exposure to airborne MPs, particularly PET fibers indoors and SBR fragments outdoors, correspond to existing literature, where textiles and tire wear are identified as major contributors to airborne MPs (Panko et al. 2019). However, discrepancies in PET fiber ratings suggest a need to review and refine data used in modeling MP fate from textiles to environmental compartments.

The model developed in this work identifies some routes with potentially high relevance for human external exposure. For LDPE, there is a high probability of external exposure by MP originating from diverse agricultural applications, such as films, packaging films, and pipes, which are finally ingested via food. However, the potential for external exposure to airborne LDPE is expected to be low. For HDPE and PP, a high probability of external exposure through ingestion via beverages and food was found for MPs originating from PCPs. In addition, the exposure to PP was also classified as high for MP originating from consumer films, bags, and packaging as well as household plastics via beverages, and agrotextiles via food consumption. SBR from tires showed high external exposure potential through food, beverages, and outdoor air, while SBR from artificial turfs only showed high results for external exposure through outdoor air. Relevant applications for PVC with the possibility of high exposure are MP released from agricultural films via food ingestion, and via ingestion of beverages for MPs from consumer films, bottles, and other packaging, as well as MPs from fabric coating. Other hot spots were identified for MPs released from consumer packaging made of PS and MPs from insulation material made of EPS via beverages. The highest exposure probability was determined for plastic applications of PET. All assessed applications received a high result for the external exposure potential via indoor air inhalation, while the potential of MP inhalation via outdoor air was assessed with a medium potential. In addition, high results were obtained for ingestion via beverages for all consumer applications, and MP exposure from agrotextiles via food was rated with high potential.

### 4.3 Translocation through the Lung and GIT

Data for the last step, involving translocation through the respiratory and the gastrointestinal tract (GIT) are limited and require further research to make polymer-specific predictions. While size barriers for translocation through tissues and cells are understood, specific data on polymer shapes and types are lacking for assessment. Due to these knowledge gaps, the likelihood of translocation of MP across tissue barriers of lung and GIT is assumed to be medium.

Our approach to estimating the human exposure potential for MP resulted in a medium and high internal exposure level for PET by ingestion and inhalation, respectively, both indoors and outdoors, while the exposure potential by dermal contact is estimated to be low (Table 9). Table 9 shows that all evaluated PET applications received a high rating for the indoor air exposure pathway. Several other applications were also found to have a high potential for ingestion via food. The exposure potential for LDPE and HDPE is classified as low for dermal contact and medium for inhalation and ingestion in most cases. Only a few LDPE and HDPE applications received a high rating for the exposure potential by ingestion. For PS, our model shows only one plastic application with a high rating, namely PS for consumer packaging via the exposure route of ingestion of beverages. Other applications were classified as medium. The results for all polymer types investigated can be found in the supporting information.

**Table 9.**
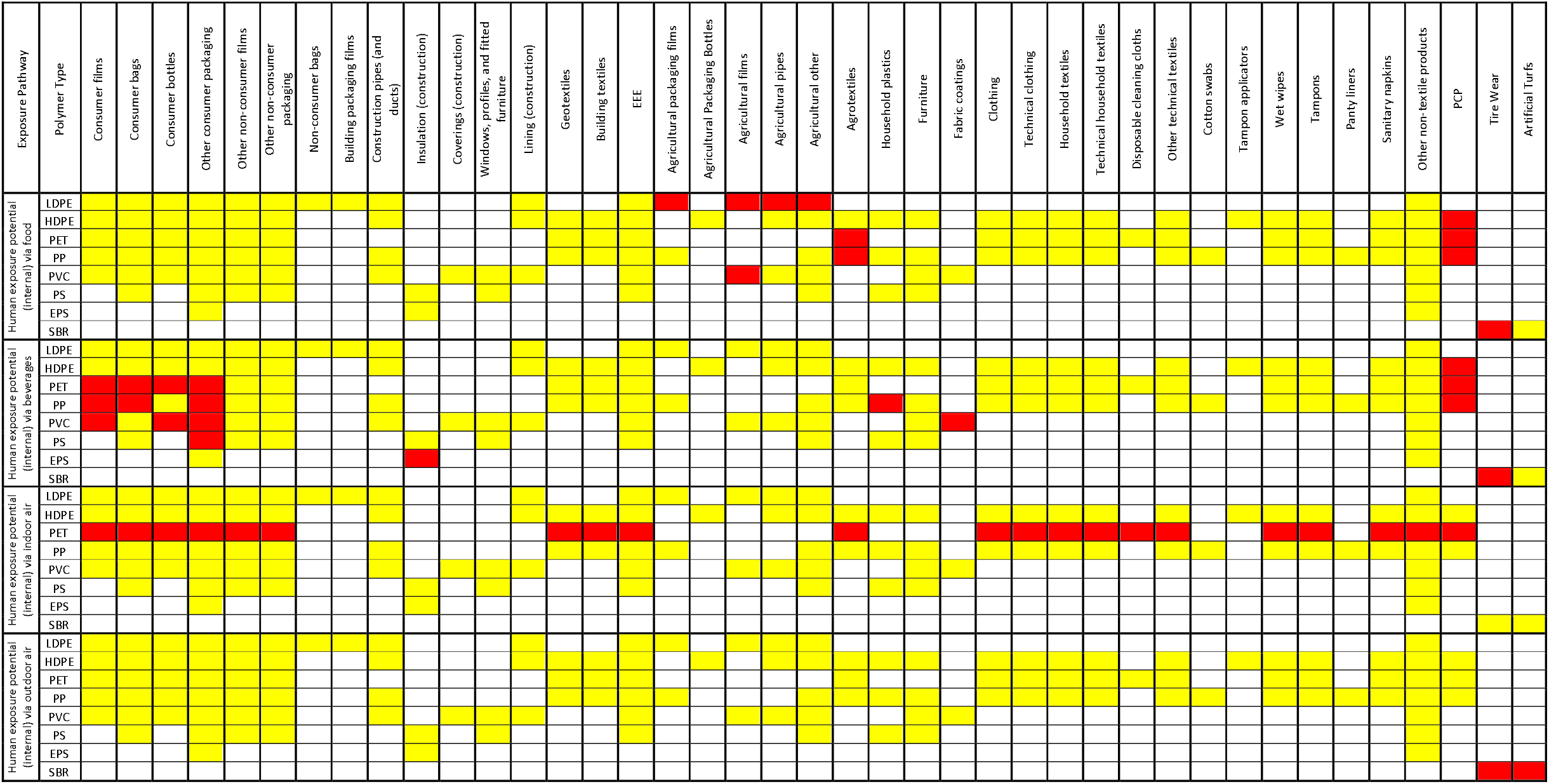
Final results for internal exposure probabilities via ingestion and inhalation for applications of LDPE, HDPE, PET, PP, PVC, PS, EPS, and SBR. Red signals high exposure, yellow medium exposure, and green low exposure. No applications have been assigned to the polymers for non-colored cells.

### 4.4 Comparison with MP Types Identified in Human Tissues

The exposure probabilities for different polymers determined using the approach presented in this paper tend to show agreement with the analysis of the occurrence of MP in humans. Yan et al. (2022) report on the frequency of polymer types of MP found in human feces. PE (as LDPE and HDPE), PET, PP, and PVC, which according to our results have more than one application with a high probability of internal human exposure via food and beverages (Table 9), are among the six most frequently found polymer types in human feces according to the results of Yan et al. (see Figure 3 in Yan et al., 2022). In the order of relative abundance according to Yan et al., PS at position 10 is the only other polymer that is also covered by our current approach. For PS, our model shows only one application that leads to a high internal exposure probability through the consumption of beverages as shown in Table 9.

Concerning exposure through inhalation, the studies by Jenner et al. (2022) and Amato-Lourenço et al. (2021), which investigated the presence of MP in lung tissue, provide an orientation as to which polymer types are to be expected in the lungs. Amato-Lourenço et al. (2021) found PP, PE, PVC, PA, polyethylene-co-polypropylene, PS, polystyrene-co-polyvinyl chloride, and PU as non-biobased MP polymers in their tissue samples. Of the polymer types covered by our approach, Jenner et al. (2022) list PP, PET, PE, and PS in descending order among the six types most frequently found in lung tissue. For PP, PE, PS, and PVC, our model predicts a medium internal exposure probability by inhalation (Table 9). For PET, the model predicts a high internal exposure probability through inhalation of indoor air. However, SBR is not included in the results of Jenner et al. and Amato-Lourenço et al. although our results show a high internal exposure probability from inhalation of outdoor air. This could be due to the limited ability to detect black tire wear particles using spectroscopic methods (Wagner et al. 2018).

In order to further verify the exposure model presented in this paper, it would be beneficial to compare it with other models. However, even if the information obtained provides only weak guidance on the validity of the models, a comparison should be made as soon as other modelling approaches for human exposure to microplastics are available that cover the entire pathway from plastic use to human exposure.

### 4.5 Limitations

The study identifies several limitations, including data gaps and the limited comparability of existing studies, which necessitated the adoption of a simplified three-class exposure model. Certain pathways, such as microplastic transport via erosion and wind, were excluded due to insufficient data, and environmental emission estimates rely on generalized values, as detailed calculations are only available for specific applications, such as textiles. The model used in the study, based on Kawecki and Nowack (2019), underestimates emissions for MPs emitting to surface water due to not including erosion and winds, and lacks differentiation for textiles and other sources. This results in oversights in critical pathways and reduces the accuracy of the findings. With regard to human exposure, the study highlights the paucity of data on polymer-specific translocation through tissues, thus rendering precise predictions challenging. Inconsistent methodologies across studies further impede comparisons, particularly with respect to food and beverage contamination levels.

Further research is thus required to address these limitations, particularly in understanding the effects of food processing and packaging on contamination, refining textile-related models, and incorporating overlooked pathways like wind and erosion into future assessments.

## 5. Conclusion

Current findings indicate widespread contamination by MPs across various environmental compartments, including the food chain, posing health risks to humans. While certain areas like MP uptake by fish and mussels have been extensively studied, others, such as contamination of food products and the influence of processing and packaging, lack thorough research. Knowledge gaps also exist in understanding degradation processes and plastic uptake by plants. Long-term studies are needed to comprehend degradation under different environments. The assessment highlights issues with data gaps and data quality particularly regarding textiles and PET fibers, impacting modelling accuracy. Rubber emissions from tires and MP ingestion through food and beverages, like vegetables and bottled water, pose significant concerns. Certain applications, like plastics in agriculture and sanitary items, require careful assessment due to limited data. Nevertheless, the results of the model represent a successful first attempt to integrate a wide range of published data on the exposure potential of plastic applications, taking into account environmental compartments, food and beverages, and uptake by humans. Further research using harmonized analytical methods is needed to allow detailed refinement of exposure prediction. A current challenge is the lack of data on the potential for uptake into the human body and on possible endpoints. Given these gaps in knowledge, it is currently not possible to make precise predictions about the risk of MP for humans. Our findings highlight the consequences of the current form of industrialization and overconsumption of plastic products. It is crucial to prevent or reduce the release of macroplastics and MPs into the environment, as these persistent materials can potentially harm organisms and ultimately also return to humans.

## Author Contributions

**Michelle Klein:** Conceptualization, Data Curation, Formal Analysis, Investigation, Methodology, Validation, Visualization, Writing - original draft, Writing – review and editing. **Bernd Giese:** Conceptualization, Funding acquisition, Methodology, Project administration, Supervision, Validation, Writing – review and editing.

All authors have read and agreed to the published version of the manuscript.

## Funding Information

This work received funding from the European Union’s Horizon 2020 Research and Innovation program, under Grant Agreement number 965367 (Project title: “Plastics fate and effects in the human body”, PlasticsFatE).

## Declaration of Competing Interest

The authors declare no competing interests.

## Notes

### Competing Interest Statement

The authors have declared no competing interest.

