## Supplementary Information for "Application-dependent assessment of the human exposure potential to microplastics"

**Supporting information**

1 Application Categories
  1.1 Consumer

Kawecki and Nowack (2019) in most cases used one Transfer Coefficient to model all polymers relevant for an application category. Apart from one other category in the application area “Building & Construction” (“insulation”), polymer specific TCs were used for “consumer film”, “consumer bags”, “consumer bottles” and “other consumer packaging”.

The definition of consumer packaging is based on a report published by WRAP (2011). The contained products include food and beverages, toiletries, cleaning products, toys and electricals.

No detailed descriptions of the plastic applications were given by either by Kawecki et al. (2018), nor by Kawecki and Nowack (2019). Therefore, the objects contained in the category “other consumer packaging” are assumed to be different from the products defined by WRAP (2011).

Kawecki and Nowack (2019) assume in their model, that consumer packaging is prone to being consumed to-go. Therefore, TCs for on-the-go-consumption are applied, which lead to littering in different environments. Following the littering in natural environment, MNPs end up in surface water. Littering in residential areas does not impact the surface water, as it is assumed, that 100% of residential litter is picked up by municipal waste handlers and ends up in the mixed waste collection.

While littering is described as a spontaneous decision, which happens in the moment, that the product reaches its end-of life, dumping is a planned act of avoidance of disposing costs (Kawecki and Nowack 2019). Products with environmental impact that are not considered in the model as being consumed to-go and end up littered, are expected to be dumped. 33% of MNPs from dumped products end up in natural environment.

1.2 Non-consumer

No detailed definition of the plastics applications of non-consumer goods are given by either by Kawecki et al. (2018), nor by Kawecki and Nowack (2019). The definition of “non-consumer” is based on WRAP (2011), which defines non- consumer plastics as plastics, which is not sold with product to consumers. This includes packaging in the retail and wholesale sector, hospitality and offices.

In contrast to consumer products, non-consumer products are not expected to be consumed on-the-go. They still have an environmental impact due to dumping and littering in residential areas. As mentioned before, 33% of MNPs from dumping end up in natural environments, which leads to the pollution of surface water. In the model by Kawecki and Nowack (2019) no air deposition from MNPs of littered products onto agricultural soil is assumed.

1.3 Building and construction

Building and Construction is category wise the biggest application area, as plastics play a major role. Here geotextiles and building textiles are considered, just as insulation, window and door frames, coverings, pipes and films.

Of these, the only category where polymer-specific TCs are considered, is the category “Insulation”. Only PS and EPS are relevant for emissions into outdoor air. All other TCs are equal for each considered polymer.

As described by Kawecki and Nowack (2019), 100 g of EPS-particles are released during the sawing and cutting per tonne EPS. As only few data are available on MNPs release during building activities and usage of building material, a TC of 0,001 was assumed for PP from pipes to soil and for wall and floor coverings to indoor air.

1.4 Agriculture

Under this aspect textiles, films, pipes, and packaging specifically used in the context of agricultural usage are considered.

According to Kawecki and Nowack (2019) the release of MNPs of plastic films is widely investigated, but shows large uncertainties. For ”agricultural films” a release of 2% as MNPs is assumed by Kawecki and Nowack (2019), ”agricultural pipes” release 0,1% as MNPs to agricultural soil.

Other than the former mentioned plastic applications, plastics used for agricultural purposes are prone to degradation processes of macroplastics due to weathering and wearing and tearing, which leads to the release of MNPs. Therefore, degradation coefficients were applied to released macroplastics in the environment.

1.5 Sanitary items

Another pathway to enter the environment is the flushing of products in toilets. This way of disposal is common for sanitary items. The sanitary items considered in this model are ”tampons”, ”tampon applicators”, ”wet wipes”, ”panty liners”, ”sanitary pads” and ”cotton swabs” (Kawecki and Nowack 2019).

Those products enter the environment through dumping. According to the TCs defined by Kawecki and Nowack (2019) 0,027% of sanitary products are dumped and fractions of this enter the surface water.

1.6 Textiles

One of the major contributors of released fibrous microplastic (FMPs) is the sector of clothing and textiles for other purposes. In 2019 synthetic fibres represented 68,4% (73,5 million tons) of global textile fibre production and further growth of the synthetic fibre market is forecasted (Garside, 2020; Global Synthetic Fibres Market Report 2020).

FMPs from clothing and textiles is released due to mechanical forces such as abrasion and aging during washing, drying and wearing (Zhang et al. 2021; Kawecki and Nowack 2019). For other textiles contributing factors are weathering, fragmentation, and dumping (Kawecki and Nowack 2019).

According to Kawecki and Nowack (2019) ”clothing”, ”technical clothing”, ”household textiles”, ”agrotextiles” and ”geotextiles” were among the top ten of the largest contributors of MP released into the water and soil. Due to lack of data no differentiation was made for PET, PP and HDPE textiles. The pathways for textile emissions that were considered in the model are washing, drying and wearing. “Household textiles” include tea towels, towels, bed linen, bedding, mattress covers, cushion covers, cushions and curtains.

Between the different types of textiles different assumptions were taken into account for the release of FMPs. While clothing textiles are assumed to undergo 19 washing cycles in the lifetime, household textile are assumed to be washed 5,9 times before being discarded.

Despite different assumptions for different textile types, generalized TCs were applied in the model by Kawecki and Nowack (2019) and in this work. 0,0027% enter the environment through “dumping”, whereby littered textiles in residential environments and road sides are fed into the mixed waste collection, while roughly 50% litter in natural environments enters surface water.

1.7 Other

Under this more general aspect categories such as ”EEE”, ”furniture”, ”household plastics” and ”PCPs” can be considered.

Kawecki and Nowack (2019) assumed a release of 0,1% to indoor air through wearing processes. Other plastic applications received a general TC of 0,00027 for the pathway to dumping and therefore resulting emissions to the environment. Other than that those product categories were neglected due to the absence of data, according to the authors.

The terms cosmetics and Personal Care Products (PCPs) are often used simultaneously, even though definitions describe essential differences.

Cosmetics, according to the European Commission, are defined as follows: “any substance or mixture intended to be placed in contact with the external parts of the human body (epidermis, hair system, nails, lips and external genital organs) or with the teeth and the mucous membranes of the oral cavity with a view exclusively or mainly to cleaning them, perfuming them, changing their appearance, protecting them, keeping them in good condition or correcting body odours.” (EU Regulation 1223/2009, Article 2.1.a) (European Union 20019).

PCPs on the other hand are not defined by law and are either classified as cosmetics or as drugs. Here cosmetics include “skin moisturizers, perfumes, lipsticks, fingernail polishes, eye and facial makeup preparations, shampoos, permanent waves, hair colours, toothpastes, and deodorants”, while the products referred to as drugs are “skin protectants (such as lip balms and diaper ointments), mouthwashes marketed with therapeutic claims, antiperspirants, and treatments for dandruff or acne.” (FDA 2022)

As in this model the most important criterion is the way of usage, specifically application to the skin, no separation between both types of products is done and the term PCPs is used.

Bans on the use of MNPs in PCPs have been enforced in several countries, including France, Sweden, United Kingdom and Italy. The definitions within these regulations are incoherent regarding the size range and a global ban and universal approach towards eliminating the use of MNPs in PCPs is still missing (Kentin and Kaarto 2018).

The most used MNP material in PCPs is PE. Depending on the function one likes to enhance by using MNPs in cosmetics, e.g. glitters, exfoliation, abrasion, viscosity regulators and so on, different shapes and sizes are used. Apart from PE other polymers such as PET and PP are used (Cole et al. 2011; Yurtsever 2019; Gouin et al. 2015).

Microbeads are used in a diversity of PCPs, such as toothpaste, shower gels, eye shadows or nail polishes, where they act as exfoliating or cleansing agents. After usage, the cosmetics and the MNPs wash out through wastewater and enter the WWTPs for treatment or get directly discarded into the environment. In a WWTP with secondary water treatment up to 95,65% of MNPs can be retained. Up to 15 billion particles per day could pass this treatment steps and enter the environment (Michielssen et al 2016).

For this model, Kawecki and Nowack (2019) consider a direct transfer of 5% of PCPs to the mixed waste collection. This is described by an average amount of 5% of products being left in the containers when being discarded (Wang et al. 2016). The other 95% of MNPs are rinsed off and enter the intermediate compartment waste water, from where it enters the process of the WWTP. The detailed TCs for the waste water treatment process are listed in the Appendix (Chapter 8.1).

1.8 Tires and artificial turfs

Tire wear particles are the largest contributor to the global MNP emissions. Even though tires are not composed of plastic but of natural and in this case most importantly synthetic rubber, rubber is an elastomer and debris should therefore be considered MNP (Kole 2017). In addition to 40-60% rubber, tires consist of carbon black, fillers, metals, textiles and additives (Evans and Evans 2006; Wagner et al. 2018). In this model, a SBR content of 50% is considered.

Wagner et al. (2018) expects soil to be heavily contaminated with debris from tire wear through runoff from roads and roadsides. Further, it depends on the local water collection and treatment system, as well as on the amount of runoff entering the aquatic environment.

Sieber et al. (2020) conducted a material flow analysis of tire wear debris in Switzerland considering three environmental sinks: roadsides, soil and surface water. Passenger cars, vans and trucks were selected as relevant vehicle categories. The data published in the study are used for this modelling approach.

The other source of SBR debris arises from artificial turfs, mainly used for soccer, rugby, golf, tennis, athletic tracks or for playgrounds (Lassen et al. 2015).

As a result of the European Landfill Directive in 2005, ELT (End-of-Life Tires) were banned from being landfilled. A new application for rubber granulate was found as infill on sports fields and playground turfs, as it is a cheap, elastic and durable alternative. 43% of ELTs are now used for artificial turfs, 21% alone on sports fields. As ELT granulates contains approximately 46% polymers they are considered MNP (Verschoor et al. 2021). The data in the model by Sieber et al. (2020) show, that from material recycling of tires 50% are used for retreading of tires and 50% are used as granulate, whereby 30% are for artificial turfs, 1% for paving, 24% for playgrounds, 24% for moulded objects and 21% for other uses.

According to a study conducted by the EU, MNP emissions from artificial turfs do not play a major role, like tire wear, textiles, paint and pellets. Emissions can be quantified with <<1% of total MNP emissions. ECHA estimates that around 10% (up to 500 kg) per field annually is lost to the environment, while the EU study suggested approximately 1500-5000 kg per field per year, with an estimated number of 13,000 large turfs in the EU in 2017, not considering a higher number of smaller pitches.

Around 100 - 120 thousand kg ELT granulate is applied on new fields to support synthetic grass fibres. During usage periodical refills an agitated zone, such as the goal and midpoint area are necessary, as the thickness of the infill layer decreases due to infill loss or compaction. Rubber granulates can also be dispersed during maintenance. Verschoor et al. 2021 quantifies annual losses as follows:

- Snow clearance: 0-150 kg
- Leaf blowing: 0-280 kg
- Brushing/Raking: 0-140 kg
- Residues on machines: 24 kg
- Players: 12-40 kg

These estimations are individual, as they depend on local and climatic conditions, seasons, methods of maintenance and awareness of players, caretakers and constructors.

| **Initial categories** | **Model categories** |
| --- | --- |
| 1. Consumer films | 1. Consumer films |
| 2. Consumer bags | 2. Consumer bags |
| 3. Consumer bottles | 3. Consumer bottles |
| 4. Other consumer packaging | 4. Other consumer packaging |
| 5.1 Other non-consumer films | 5. Non-consumer films, bags, packaging |
| 5.2 Other non-consumer packaging |  |
| 5.3 Non-consumer bags |  |
| 6. Building packaging films | 6. Building packaging films |
| 7. Construction pipes (and ducts) | 7. Construction pipes |
| 8. Insulation (construction) | 8. Insulation (construction) |
| 9. Coverings (construction) | 9. Coverings (construction) |
| 10.1 Windows, profiles and fitted furniture | 10. Windows, profiles, lining, fitted furniture |
| 10.2 Lining (construction) |  |
| 11. Geotextiles | 11. Geotextiles |
| 12. Building textiles | 12. Building textiles |
| 13. EEE | 13. EEE |
| 14.1 Agricultural packaging films | 14. Agr. Packaging |
| 14.2 AgriculturalPackagingBottles |  |
| 15. Agricultural films | 15. Agr. Films |
| 16.1 Agricultural pipes | 16. Agr. Pipes and other |
| 16.2 Agricultural other |  |
| 17. Agrotextiles | 17. Agrotextiles |
| 18. Household plastics | 18. Household plastics |
| 19. Furniture | 19. Furniture |
| 20. Fabric coatings | 20. Fabric coatings |
| 21.1 Clothing | 21. Different textiles |
| 21.2 Technical clothing |  |
| 21.3 Household textiles |  |
| 21.4 Technical household textiles |  |
| 21.5 Disposable cleaning cloths |  |
| 21.6 Other technical textiles |  |
| 22.1 Cotton swabs | 22. Sanitary items and other non-textile products |
| 22.2 Tampon applicators |  |
| 22.3 Wet wipes |  |
| 22.4 Tampons |  |
| 22.5 Panty liners |  |
| 22.6 Sanitary napkins |  |
| 22.7 Other non-textile products |  |
| 23. PCP (Personal Care Products) | 23. PCP |
| 24. Tire | 24. Tire |
| 25. Artificial Turfs | 25. Artificial Turfs |

2 Land Use Data

The models used to estimate the share of plastics in environmental compartments referenced Switzerland (Sieber et al. 2021; Kawecki and Nowack 2019). Therefore, adjustments were made to align calculations with EU conditions. Land-cover data from Eurostat (2021) were incorporated, modifying percentages of surface water, residential soil, agricultural soil, and natural soil. For instance, the EU has 3,2% surface water compared to Switzerland's 4,3%. Similarly, residential soil accounts for 4,2% in the EU compared to 7,5% in Switzerland. Agricultural soil in the EU is 24,2% compared to Switzerland's 35,9%, while natural soil is estimated at 68,3% for the EU and 52,3% for Switzerland. These adjustments were integrated into the normalized model.

| Classifications by Eurostat | Land-cover in % | Aggregation, value in % of total landcover | Classification by Kawecki and Nowack (2019) and Sieber et al. (2020) | Landcover in % in Kawecki and Nowack (2019) and Sieber et al. (2020) |
| --- | --- | --- | --- | --- |
| Water Areas | 3,2 | 3,2 | Surface Water | 4,3 |
| Artificial Land | 4,2 | 4,2 | Residential Soil | 7,5 |
| Cropland | 24,2 | 24,2 | Agricultural Soil | 35,9 |
| Wetland | 1,7 | 68,3 | Natural Soil | 52,3 |
| Bareland | 2,4 |  |  |  |
| Shrubland | 5,7 |  |  |  |
| Grassland | 17,4 |  |  |  |
| Woodland | 41,1 |  |  |  |

3 Food and Beverages
  3.1 Meat

Kedzierski et al. (2020) focuses on MNP contamination of chicken meat in PS packaging. In order to determine the amount of fibres and particles on the outside of the meat, it was rinsed with distilled water, which was then collected and further analysed. Additionally, the inside of the packaging was rinsed and the water with the retrieved particles collected. The reported amount is 4-19 MNPs/kg of packaged meat.

The per capita consumption in the EU of poultry is stated to be 22,21 kg in 2019, which resonated to an annual consumption of 88,84 – 421,99 MNPs/year and person (Ritchie et al. 20117). As a pessimistic and careful approach is taken, the ultimate value taken for the assessment is 422 MNPs/year.

It is highly likely that the contamination occurred through PS dust being suspended in the air of the production site of the packaging trays, therefore before the packaging of the meat products. The particles are easily airborne due to their low mass and size, and because of the electrostatic properties of PS make them stick to the surface. This theory also resonates with the fact, that MNP particles were found between the meat and the plastic film of the packaging (Kedzierski et al. 2020).

Another publication focused on the influence of plastic cutting board as a source of MNPs in meat products (Habib et al. 2022). The cutting boards were made of PE and all meat samples showed debris identical with the material of the boards. On average 1,19 ± 0,41 mg of MP was found per g of unwashed meat. Another finding concluded that older cutting boards release more MP than newer ones. When washed for 10 seconds, the content of MP in the meat was only insignificantly reduced. Extending washing time to three minutes lead to a reduction of MP content from up to 97 MNPs to approximately 1 piece per 15 g of meat (Habib et al. 2022).

Based on these results, Habib et al. (2022) calculates up to 70 MP/kg of red meat even after washing for 3 minutes. Assuming a yearly consumption of 20,46 kg of red meat, this sums up to an annual intake of 1432,2 MNP particles per year (Salter 2018).

3.2 Fish and Seafood

MNPs in fish and seafood have been widely investigated as marine areas are under increasing pressure by human activities and pollution. 80 to 85% of marine litter is assumed to be plastic (Auta, Emenike et al. 2017). MNPs enter the marine environment through several pathways and from various applications, such as toothpaste, resin pellets, facial scrubs and plastics that undergo processes of fragmentation. Common pathways are through domestic and industrial drainage systems and wastewater treatment plants (WWTPs), another source are waste dumps (Cole et al. 2011; Murphy et al. 2016; Alomar et al. 2016). Due to their small size, MNPs are easily available for ingestion by marine organisms such as bivalves, mussels, fishes, shrimps and oysters, which are also relevant for the human food chain (Cole et al. 2013).

Shellfish, Squid and Shrimp from the Arabian Sea were investigated by Daniel et al. (2021).

From shellfish, an average of 2,7 MNPs/kg edible tissue were extracted. This sums up to 0,07 pieces per individual shellfish. The results for squid were higher, with 0,18 particles per individual and 7.7 MNPs/kg edible tissue. For crab meat the results were 0,14 MNPs/individual and 3,2 MNPs per kg of edible issue. PP was the most prominent polymer with 40%, followed by PE with 27% and PS with 20%.

According to a report by the FAO (2016) the global per capita annual consumption of shellfish is estimated to be 4,9 kg. This would indicate that approximately 13 pieces of MNP are ingested annually per person by eating shellfish.

A recent study by Gündoǧdu and Köșker (2023) examined a diverse selection of different canned fish sold in Türkiye. The samples included canned Black Sea anchovy, Norwegian salmon, longtail tuna, yellowfin tuna, skipjack, and mackerel fish. Some samples additionally contained olive or sunflower oil and salt, water, and lemon water. While differences in fish species, packaging types and the additional salt and oil had no significant impact on the MNP content, Gündoǧdu and Köșker found a significant difference depending on the producer.

Packaging presumably contributed to contamination by MNP to the canned fish. Can manufacturers started using acrylic, polyolefins and polyester as coatings instead of epoxy due to public concerns. This is represented in the results, as the majority of found MNP in these samples consisted of those polymers used for can coating.

The samples contained between 0,81 and 13,50 particles per 100g of fish flesh, resulting in a mean across all samples of 4,12 particles/100 g. The share of fragment shaped particles was estimated at 57,4% and fibres represented 42,7%.

According to the European Commission (2021) the annual consumption of fish in the EU is 23,97 kg. With a contamination rate 4,12 MNPS/100g the annual uptake sums up to 988 particles in total, including polymer types which are not relevant for this work.

3.3 Fruit and vegetables

In a 2023 study, Aydin et al. investigated the presence of MNP debris in different commonly consumed fruits and vegetables, specifically in apples, pears, tomato, onion, potato, and cucumber. The number of particles was quantified, and an EDI (Estimated Daily Intake) was calculated. The samples were washed, peeled, and sliced.

The absorption and translocation into plant tissues is assumed to occur similar to the process of carbon nanomaterials (Herremans et al. 2015). The uptake by the plants system is inversely proportional to its size. Larger particles are unable to penetrate into the plants system and therefore get adsorbed on the surface, while smaller particles get absorbed through the roots or in seeds through endocytosis and translocate into the shoot and from there through capillary action into the aerial parts (Husen and Siddiqi 2014).

The vascular system of fruits is more pronounced than the vascular system of vegetables, because of necessary nourishment and protection of the seeds. This results in a greater accumulation of MNPs in fruits due to a higher vascularization of the fruit pulp. Additionally, the root system of fruit trees is larger than that of vegetables and the age of the fruit tree with a life span of several years compared to a life time of several days of a carrot is more extensive (Oliveri Conti et al. 2020).

3.4 Rice

A study examining plastic content in rice samples was carried out in Australia. Different brands and types of rice were bought in stores. This included uncooked rice in HDPE plastic bags, instant rice in PP plastic bags, uncooked rice in a cloth bag as well as uncooked and unpacked rice from a bulk store, which was then refilled into paper bags (Dessi et al. 2021).

PE was detected in all rice samples with a ranging concentration from 45 – 317 μg/g of rice. In 40% of the samples PP was found with a maximum concentration of 105 μg/g and 6% of the samples contained PET with a concentration of 17 μg/g at the maximum. The results thereby varied by the packaging material, the type of rice (instant rice and uncooked rice) and the form of treatment during testing. Some samples were washed and/or shaken. Notably, PET was only present in instant rice and could be completely removed by washing. Instant rice also had an overall higher concentration of PE than the other types of rice (Dessi et al. 2021).

Other factors as origin of rice, farming techniques and qualities of rice had no impact on the MNP contamination of the rice. Washing reduced the MNP content of around 20% in uncooked rice and 40% in instant rice. Shaking of the package was assumed to increase the content of MNP due to abrasion, but that assumption was disproved (Dessi et al. 2021).

Assuming a 100g portion of cooked rice as serving unit, the intake of MNP for unwashed rice sums up to 3,7 mg, for washed rice 2,8 mg and for instant rice 13,3 mg (Dessi et al. 2021).

The annual consumption of rice in the EU is 6,04 kg (OECD-FAO 2021).

As no size and shape for the MNPs was measured, an assumption needs to be made in order to calculate a number of MNPs in the samples. Those calculations are available in the Supporting Information.

3.5 Honey

Contamination of honey has been thoroughly investigated by Liebezeit (2015). 10 - 336 fibres/kg of undefined polymer type were found. The amount of fragments was lower, with 2 - 82 fragments/kg. All 47 honey samples were contaminated with MNP fibres and particles. The plastic particles are airborne and result from various applications, abrasion processes and deposit on flowers, which is then transported to the hive by insects (Liebezeit 2015).

Another sampling of honey was done by Diaz-Basantes et al. (2020), were industrial honey and craft honey were tested for MNP particles. Here the industrial honey contained 20-166 fibres/ and 126 - 552 fragments/litre. The craft honey showed contamination of MNP debris in a similar range, with 82-178 fibres/litre and 200 - 828 fragments/litre.

When assuming an equal consumption of the EU to the US, 0,61kg are annually consumed per capita (Soocial 2022). Based on data provided by Diaz-Basantes et al. (2020) this would lead to the following annual consumption of MNPs:

3.6 Salt

Four different types of salt can be distinguished: sea salt, lake salt, rock salt and well salt. While samples collected in Asia show a higher contamination rate, also a difference between different salt types was found. The pollution of salt samples was found to depend on the pollution of the aquatic system in several studies (Jin et al. 2021).

Samples from different European locations were investigated, including France, Italy, UK, Germany, Bulgaria (produced in Belarus), Hungary (produced in Romania) and Croatia. All samples were sea salt, except for samples from Germany, Italy, Belarus and Romania which are rock salt. The share of polymers found in sea salt were identified as 35% PE, 30% PP, 30% PET, the ones of rock salt as 26% PE, 23% PP, 41% PET. Sea salt contained 0-1674 MNPs/kg and rock salt 0-148 MNPs/kg. In sea salt 63% of MNPs were described as fragments and 31% as fibres. For rock salt the share was 54% and 45% respectively. A third shape was identified as sheets, which amounted to 6% and 1% (Kim et al. 2018).

The salt intake varies between different countries and regions and also between men and women, as men consume more salt. With an average of 9,95 g daily intake of salt, the annual consumption sums up to 3,63 kg per capita (Kloss et al. 2015).

When taking a pessimistic approach and assuming a content of 1674 MNPs/kg sea salt, the annual uptake sums up to 6.077 MNPs per year. For rock salt the annual uptake is 537 MNPs/per capita/year.

3.7 Sugar

Only few sugar samples have been investigated so far, but all of them contained MNPs. Liebezeit (2013) selected two samples of refined sugar and two samples of powdered sugar. Additionally one sort of unrefined cane sugar was sampled. The measured amount of found MNPs in the samples sums up to 217 ± 123 fibres/kg and 32 ± 7 fragments/kg. The cane sugar contained 560 fibres and 540 fragments/kg (Liebezeit 2013).

The OECD/FAO (2021) calculated a per capita sugar consumption in the EU of 37,6 kg. For refined and powdered sugar the annual consumption adds up to 8159 fibres and 1203 fragments, therefore 9362 MNPs in total. For cane sugar the annual per capita consumption 21056 fibres and 20304 fragments, a total of 41360 MNPs.

3.8 Tap water

Several studies examined the MNP concentration in tap water, but only two were available with data from Europe, as most focused on Asia.

Johnson et al. (2020) focused on tap water in England and Wales from different sources. The average contamination rate of raw water from the water treatment facility showed a contamination rate of 4,9 MNPs/L.

39 samples of potable water were compared and overall showed low contamination rates. Only one particle of PS and two particles of ABS were quantified in a volume of 14,2 m³ of water for one sample.

The average values of MNP pollution in raw and potable water were calculated to 4,9 MNP/L and 0,00011 MNPs/L respectively. This indicates an extremely effective filter process at the water treatment facility. Overall, these low numbers resonate with results published by Mintening et al. (2018), who quantified MNP particles in 24 German groundwater samples, which were taken at different stages of the purification process (Johnson et al. 2020).

Mintening et al. (2018) detected five different polymers in the samples of which 62% were polyester, PVC contributed 14%, and PE 6%. The other polymers are not of interest for this work. The particle sizes ranged from 50 – 150 μm. MNP concentrations ranged from 0 – 7 MNPs/m³ and 14 of 24 samples contained no MNPs. In conclusion, water from groundwater shows a low contamination rate of 0,7 MNPs/m³.

No data are available on the per capita consumption of tap water for drinking purposes only, therefore a consumption rate for bottled water is applied for the calculation of the annual consumption of water and thereby MNPs. According to Statista (2023) in 2021 110,3 litre of bottled water were consumed per person per year. Applying this consumption rate to the data of Johnson et al. (2020) this would mean an annual MNP consumption of 0,01. As ABS is not considered in this work and only one third of the particles is PS, this would lead to an annual consumption of 0,004 PS particles.

According to Mintening et al. (2018) 1000 litre of drinkable groundwater contain 0,7 MNPs. For the annual consumption of 110,3 litre this would add up to 0,08 MNPs, of which 0,05 are polyester which can be considered PET-fibres, 0,01 are PVC and 0,005 are PE.

3.9 Bottled water

Bottled water was been widely investigated. Oßmann et al. (2018) selected three different packaging types of bottled water and compared plastic content, size of debris and polymer typ. The study was conducted in Germany and 21 different brands were sampled. New and older reusable bottles were distinguished by the amount of scratches and the cloudiness of the material.

The highest mean value of MNP particles per litre was found in old reusable PET bottles with a value of 8339 particles/l, while newish reusable bottles contained 2689 particles/l. Single use PET bottles have shown a similar value of 2649 particles/l and in glass bottles 6292 particles/l were found.

When applying the annual consumption rate of 110,3 litres of bottled water per year based on Statista (2021), the values and the share of polymers for the described packaging variations can be calculated.

3.10 Tea

One study by Hernandez et al. (2019) investigated the release of micro- and nanosized plastic particles of teabags in the process of tea brewing.

Four samples of individually packed tea-bags were bought in store and emptied of the dried tea leaves. This was done in order to obtain MP released by the teabags itself and not through contamination of the tea products contained in the teabags. Two samples consisted of nylon, while the other two were made of PET. As nylon is not in the scope of interest of the present study, only the results of PET will further be discussed.

Considering the size of the particles and the count of pieces per cup of tea, it is estimated that 13-16 μg of debris are consumed with each cup of tea. Converting this into a number of consumed particles, Hernandez et al. (2019) reported 2.3 million particles with a size over 1 μm and 14.7 billion particles sized smaller than 1 μm to be ingested when drinking one cup of tea using one teabag.

The European consumption rate for tea is 0,84 kg according to Statista (2023). Assuming a filling volume of 2 g per teabag (The Tea Detective 2022). The annual consumption is calculated to 420 teabags. The assumption also includes, that tea is exclusively consumed in teabags.

With a total of 14.7 billion MNPs consumed per cup of tea, the annual uptake is 6172.5 billion PET particles.

3.11 Beer

MNP contamination of beer was investigated by Diaz-Basantes et al. (2020). The selected samples contained on the one hand industrial beer and on the other hand craft beer filled in glass bottles. Beer in glass bottles was preferably selected as to eliminate the packaging as a source of contamination. As beer is mainly made of water, this also poses as the prime source of MNP contamination.

Industrial beer shows a higher contamination rate than craft beer with 47 MNPs/L and respectively 32 MNPs/L.

According to data published by Brewers of Europe (2022) the average consumption in European countries is 63,2 litres per capita per year. For industrial beer this would result in an annual consumption of 2970 MNPs and for craft beer the uptake would sum up to 2022 MNPs.

3.12 Skim milk

The MNP content of skim milk was researched by Diaz-Basantes et al. (2020). The specific characteristics included a fat content of <1% and a packaging with plastic content - either beverage cartons or polyethylene cover.

The number of fragments and fibres counts up to 40 particles per Litre. No differentiation between polymer types was published.

The average citizen within the EU consumes 56,7 litres annually. Therefore, the intake of MNP particles via consumption of milk can be quantified with 2268 particles/year.

3.13 Softdrinks

A diverse range of soft drinks was sampled by Diaz-Basantes et al. (2020) and MNP contamination was quantified. The chosen samples are characterized as refreshing beverages with citrus flavour and were packed in either beverage cartons or PET bottles. The presence of a different polymer than those used as packaging material directly indicate a source of contamination during the production process.

The detected amount of found particles resembles those of craft beer, with 32 particles/litre.

To calculate to daily intake based on consumption, data provided by the UNESDA (2023) – the European soft drinks industry – were used. They reported a per capita consumption of 92,2 litres/year in 2021. An intake of 2950 particles/year can be calculated, based on the findings of Diaz-Basantes et al. (2020).

4 Outdoor Air and Inhalation

For tire wear particles consisting of styrene-butadiene rubber (SBR), a density of 1g/cm³ and a spheric shape is assumed. The total number of particles can be calculated from the given results (Spektrum 1998, Panko et al. 2019). The assessment of road wear particles in outdoor air by Panko et al. (2019) yielded an average of 18,35 μg/m³ of PM2,5 particles at the sampling sites in London. The amount of PM10 particles summed up to 47,43 μg/m³. For the calculation of PM10 particles the calculation is as follows:

𝑉𝑜𝑙𝑢𝑚𝑒=($\frac{4}{3}$)∗𝜋∗(5μ𝑚) ³=5,236μ𝑚³

𝑀𝑎𝑠𝑠=𝐷𝑒𝑛𝑠𝑖𝑡𝑦∗𝑉𝑜𝑙𝑢𝑚𝑒=1𝑔/𝑐𝑚³∗5,236μ𝑚³=5,236 μ𝑔

𝑇𝑜𝑡𝑎𝑙 𝑚𝑎𝑠𝑠/𝐼𝑛𝑑𝑖𝑣𝑖𝑑𝑢𝑎𝑙 𝑝𝑎𝑟𝑡𝑖𝑐𝑙𝑒 𝑚𝑎𝑠𝑠= $\frac{47,43 \mu g}{5,236\mu g}$=9,054

A measured mass of 47,43μg/m³ therefore contains approximately 9,054 SBR particles.

The same calculation is done for the PM2,5 particles, resulting in an estimated amount of approximately 2,241 SBR particles/m³ as shown in the following calculation:

𝑉𝑜𝑙𝑢𝑚𝑒=($\frac{4}{3}$)∗𝜋∗(1,25μ𝑚) ³=8,18μ𝑚³

𝑀𝑎𝑠𝑠=𝐷𝑒𝑛𝑠𝑖𝑡𝑦∗𝑉𝑜𝑙𝑢𝑚𝑒=1𝑔/𝑐𝑚³∗8,18μ𝑚³=8,18 μ𝑔

𝑇𝑜𝑡𝑎𝑙 𝑚𝑎𝑠𝑠/𝐼𝑛𝑑𝑖𝑣𝑖𝑑𝑢𝑎𝑙 𝑝𝑎𝑟𝑡𝑖𝑐𝑙𝑒 𝑚𝑎𝑠𝑠= $\frac{18,35 \mu g}{8,18\mu g}$=2,241

With 6,25m³ of daily inhaled outdoor air, a number of up to 57 PM10 and 14 PM2,5 SBR particles per day can be expected to be contained in the inhaled air.

As no size and shape for the other MNPs was measured, an assumption needs to be made in order to calculate a number of MNPs in the samples. According to definition (Thompson et al. 2004, Gigault et al., 2018; Mitrano and Wohlleben, 2020), microplastics are sized between 1µm to 5 mm, therefore we assume a spheric medium particle size of 0,25mm. The density of PE is 0,9 g/cm³ and for PP of 0,91g/cm³ (Omnexus n.d.).

The calculation for the mass of a single PE particle is as follows:

𝑉𝑜𝑙𝑢𝑚𝑒=($\frac{4}{3}$)∗𝜋∗(0,125𝑚𝑚)³=0,00818𝑚𝑚³

𝑀𝑎𝑠𝑠=𝐷𝑒𝑛𝑠𝑖𝑡𝑦∗𝑉𝑜𝑙𝑢𝑚𝑒=0,9𝑔/𝑐𝑚³∗0,00818𝑚𝑚³=7,36 µ𝑔

For PP the following calculation applies:

𝑉𝑜𝑙𝑢𝑚𝑒=($\frac{4}{3}$)∗𝜋∗(0,125𝑚𝑚)³=0,00818𝑚𝑚³

𝑀𝑎𝑠𝑠=𝐷𝑒𝑛𝑠𝑖𝑡𝑦∗𝑉𝑜𝑙𝑢𝑚𝑒=0,91𝑔/𝑐𝑚³∗0,00818𝑚𝑚³=7,44 µ𝑔

In order to obtain the contained particle amount, the mass of MNPs as described are divided by the mass of a single particle, as calculated.
